# Interplay between p300 and HDAC1 regulate acetylation and stability of Api5 to regulate cell proliferation

**DOI:** 10.1101/2020.11.22.393256

**Authors:** Virender Kumar Sharma, Mayurika Lahiri

## Abstract

Api5, is a known anti-apoptotic and nuclear protein that is responsible for inhibiting cell death in serum-starved conditions. The only known post-translational modification of Api5 is acetylation at lysine 251 (K251). K251 acetylation of Api5 is responsible for maintaining its stability while de-acetylated form of Api5 is unstable. This study aimed to find out the enzymes regulating acetylation and deacetylation of Api5 and the effect of acetylation on its function. Our studies suggest that acetylation of Api5 at lysine 251 is mediated by p300 histone acetyltransferase while de-acetylation is carried out by HDAC1. Inhibition of acetylation by p300 leads to reduction in Api5 levels while inhibition of deacetylation by HDAC1 results in increased levels of Api5. This dynamic switch between acetylation and deacetylation regulate the localization of Api5 in the cell. This study also demonstrates that the regulation of acetylation and deacetylation of Api5 is an essential factor for the progression of the cell cycle.

## Introduction

Apoptosis inhibitor 5 (Api5) is also known as anti-apoptotic clone 11, AAC-11. Api5 was discovered as a nuclear protein, which was responsible for inhibition of apoptosis under serum starvation conditions [1]. Api5 is a right handed superhelix which is composed of all helical repeats. It has 19 α-helices and two 3_10_ helices [2]. It has also been observed that all these helices interact with neighbouring helices to form antiparallel helix pair. N-terminal of Api5 comprises of α-helix from 1 to 11 which shares homology with the HEAT repeat structures of other proteins such as Importin β. C- terminal of Api5 is made up of α-helices from residues 12 - 19 and share homology with ARM-like repeat structures of other proteins, for example, p120 catenin. The N-terminal of Api5 contains LxxLL motif which is an amphipathic α-helix that forms nuclear receptor co-factor interaction regions. This motif is present in the sixth α–helix and is required for maintaining stability of the protein by forming hydrophobic interactions with neighbouring α-helices and is also the region that interacts with FGF2 [3]. The C-terminal region of Api5 has a nuclear localization signal and a putative Leucine Zipper domain (LZD). LZD is present in the α18 helix, between 371-391 amino acid residues. It interacts with Acinus, a protein which mediates fragmentation of chromatin during apoptosis [4]. Global mass spectrometric analysis of Api5 suggests the presence of one conserved acetylation site at lysine 251, which is present in the hinge region between α13 and α14 [2].

It has been proven that Api5 inhibits the dE2F1-mediated apoptosis in *Drosophila* cells [5]. dE2F1 is a well-known pro-apoptotic gene responsible for apoptosis in the fruit flies. In the *Drosophila* embryonic cells, SL2, RNAi-mediated depletion of Api5 was found to be responsible for increased apoptotic cell death as compared to the dE2F1 over-expressed cells. Similar result was also observed in the human osteosarcoma cell line, Saos-2, where ectopic expression of Api5 decreased E2F1-mediated apoptosis in E2F1 over-expressing cells without affecting its transcriptional activity [5].

According to the study by Rigou et al., AAC-11 binds to Acinus, a nuclear protein and plays a significant role in chromatin condensation during apoptosis [4]. Binding of AAC-11 to Acinus does not allow its cleavage by caspase-3, thus in turn inhibiting DNA fragmentation and apoptosis [4].

Studies performed in melanoma cells, showed Api5 to modulate FGF2 and FGFR1 signalling which activates ERK. This activated ERK phosphorylates Bim, a pro-apoptotic protein. Phosphorylated Bim is the target for proteosomal degradation. Thus ubiquitin-mediated degradation of Bim is a means by which Api5 inhibits apoptosis in HeLa and 565mel cell lines [6].

Mayank and group have reported Api5 to inhibit transcription of APAF1 gene. APAF1 is the main component of the apoptosome complex. Thus Api5 prevents the formation of the apoptosome, in turn inhibiting apoptosis [7]. A recent study suggests Api5 to physically interact with caspase 2, and prevent its activation, thus inhibiting apoptosis [8].

Api5 has also been reported to be involved in the regulation of E2F1, a transcriptional activator of G1/S cell cycle transition genes, by enhancing the binding affinity of E2F1 to its target promoters. Knockdown of Api5 arrested H1299 cells at the G1 phase of the cell cycle, thus proving that apart from regulating apoptosis, Api5 also plays a critical role in maintaining normal cell cycle progression [9].

Navarro and co-workers have demonstrated that the levels of Api5 are regulated in a cyclic manner [9]. It was observed that the levels of Api5 was higher in the G1 phase and was stabilized during the G1/S transition. Interestingly, Api5 levels decreased as the cells proceeded further in the cell cycle, from G2 to G2/M phase [9]. This suggests that Api5 undergoes degradation during cell cycle progression. Knockdown of Api5 arrests H1299 non-small cell lung carcinoma cells at the G1 phase. This was further supported by Han and his group where they showed acetylation of Api5 at lysine 251 to be associated with its stability [2].

Api5 has been found to be overexpressed in different types of cancers like cervical, urinary bladder, lung, ovarian and oesophageal cancers [6; 10; 11; 12; 13]. In cervical cancers, Api5 overexpression has been shown to promote invasion [12]. Api5 has also been shown to promote the degradation of Bim, a pro-apoptotic protein [6]. In osteosarcomas, studies have shown Api5 to inhibit E2F1 as well as Acinus-mediated apoptosis [4; 5]. Api5 has been identified as a biomarker for cervical and ovarian cancers and a prognosis marker for non-small cell lung carcinomas [11; 13]. High levels of Api5 provide cancer cells the ability to evade immune response mediated cell death [6]. In breast cancers, Api5 interacts with the estrogen receptor to promote proliferation [14]. It has also been reported that Api5 promotes metastasis in breast cancers [14]. Higher levels of Api5 are associated with chemo-resistance [15]. It has been shown that tamoxifen-resistant breast cancer cells show an upregulation of Api5 [16], while cancer cells which are sensitive to anticancer agents like tocotreinol show reduced levels of Api5 [17]. Reduced and low levels of Api5 are associated with the increase in cell death in various cancers. Knockdown of Api5 resulted in the reduction in *in vivo* tumorogenicity in cancer cells [14].

Api5 acetylation at lysine 251 is conserved from protists to mammals [2]. De-acetylated form of Api5 is not stable and therefore undergoes post translational degradation. However the mechanism of degradation and the enzymes involved in the process of acetylation and de-acetylation of Api5 is not yet known.

CBP/p300, GCN5/PCAF and TIP60/MYST1/2/3/4 are the major acetyltransferases involved in acetylation of most of the cellular proteins. Among this, p300 acetylates proteins involved in a number of diverse biological functions including proliferation, cell cycle regulation, apoptosis, differentiation and DNA damage response [18; 19; 20; 21]. p300 histone acetyl transferase was initially identified as a transcriptional activator that performs its function by acetylating histones in eukaryotic cells. p300 is capable of acetylating all the four histones [22; 23]. Later it was discovered that p300 also acetylates non-histone proteins like E2F1, p53, p73, Rb, E2F, myb, myoD, HMG(I)Y, GATA1 and α-importin [24; 25; 26; 27; 28; 29; 30; 31; 32; 33]. The role of p300 histone acetyl transferase in the regulation of the cell cycle is also known [34]. Activity of p300 is required for normal transition from G1 to S phase of cell cycle [35]. Mutations or abnormality in function of p300 leads to multiple disorders and cancer is most common amongst them [36; 37; 38; 39].

Histone de-acetylases (HDACs) are the major de-acetylases involved in the de-acetylation of histones that regulate genetic expression of certain genes as well as non-histone proteins that regulate the processes responsible for maintaining cellular homeostasis [40; 41; 42; 43]. HDAC de-acetylates proteins involved in almost all the cellular events like cell cycle, replication, proliferation and apoptosis. It has also been shown that most of the proteins that get acetylated by p300 histone acetyltransferase also undergo de-acetylation by HDAC1, a member of class1 HDACs, for example p53 and E2F1 [28; 42]. Han and group in their studies have suggested the possibility of involvement of histone de-acetylases in the de-acetylation of Api5 [2].

AMP-activated protein kinase (AMPK) is an energy sensor kinase that regulates almost all cellular processes like cell division, DNA replication, transcription activation and apoptosis [44; 45; 46] by sensing AMP to ATP ratio and thereby regulating cellular homeostasis. It has also been reported that AMPK regulates G1/S transition of cells [47]. Mutations in AMPK can lead to multiple disorders including cancer [48].

Protein kinase B (Akt) has various functions in cells that include proliferation, migration and transcription. Studies have shown PI3K/Akt pathway to also regulate cell cycle progression by phosphorylating the inhibitors of cyclin-dependent kinases (CKI) p21 and p27 Cip/Kip proteins [49; 50; 51]. Akt has also been shown to play a role in regulating different pro-apoptotic molecules in order to enhance survival and proliferation of cancer cells [52].

In this study, we report two novel and key regulators of Api5: p300 and HDAC-1. We observed p300 to be the enzyme that interacts with and regulate Api5 levels in cells. p300 mediated acetylation of Api5 at lysine 251 provided stability to Api5, while the HDAC1-mediated deacetylation led to reduced levels of Api5 in the cells. Both these regulators also regulated the subcellular localization of Api5 in cells. We observed that the de-acetylated protein was transported to the cytoplasm while the acetylated protein was present in the nucleus. The regulation of acetylation was also observed to regulate cell cycle progression. We also demonstrated AMPK and Akt to be the players regulating p300 activity, thereby leading to stability of Api5 in cells.

## Results

### *in silico* analysis predicts p300 histone acetyltransferase to acetylate Api5 at lysine 251

It has been reported in previous studies that Api5 undergoes acetylation at lysine 251, however, the enzyme that is required for the acetylation function has not yet been identified. Therefore, to identify the enzyme involved in the acetylation of Api5 we performed *in silico* analysis using the ASEB web server tool [53; 54] to predict the site as well as the enzyme which may be responsible for acetylation of Api5 at lysine 251. It was observed that amongst CBP/p300, GCN5/PCAF and TIP60/MYST1/2/3/4, CBP/ p300 had a lower p value suggesting a higher possibility of acetylating Api5 at lysine 251 (Figure 1A). Active site residues of p300 histone acetyltransferase were obtained from the literature [55]. Docking was performed between p300 and Api5 using the HADDOCK webserver [56]. The output was analyzed using Protein Interaction Z Score Assessment (PIZSA) [57] and the best scoring model was chosen as shown in figure (1B-D). Interestingly the acetylating domain of p300 showed interaction with three amino acid residues of Api5 including lysine 251 (Figure 1C-D).

**Figure 1.**
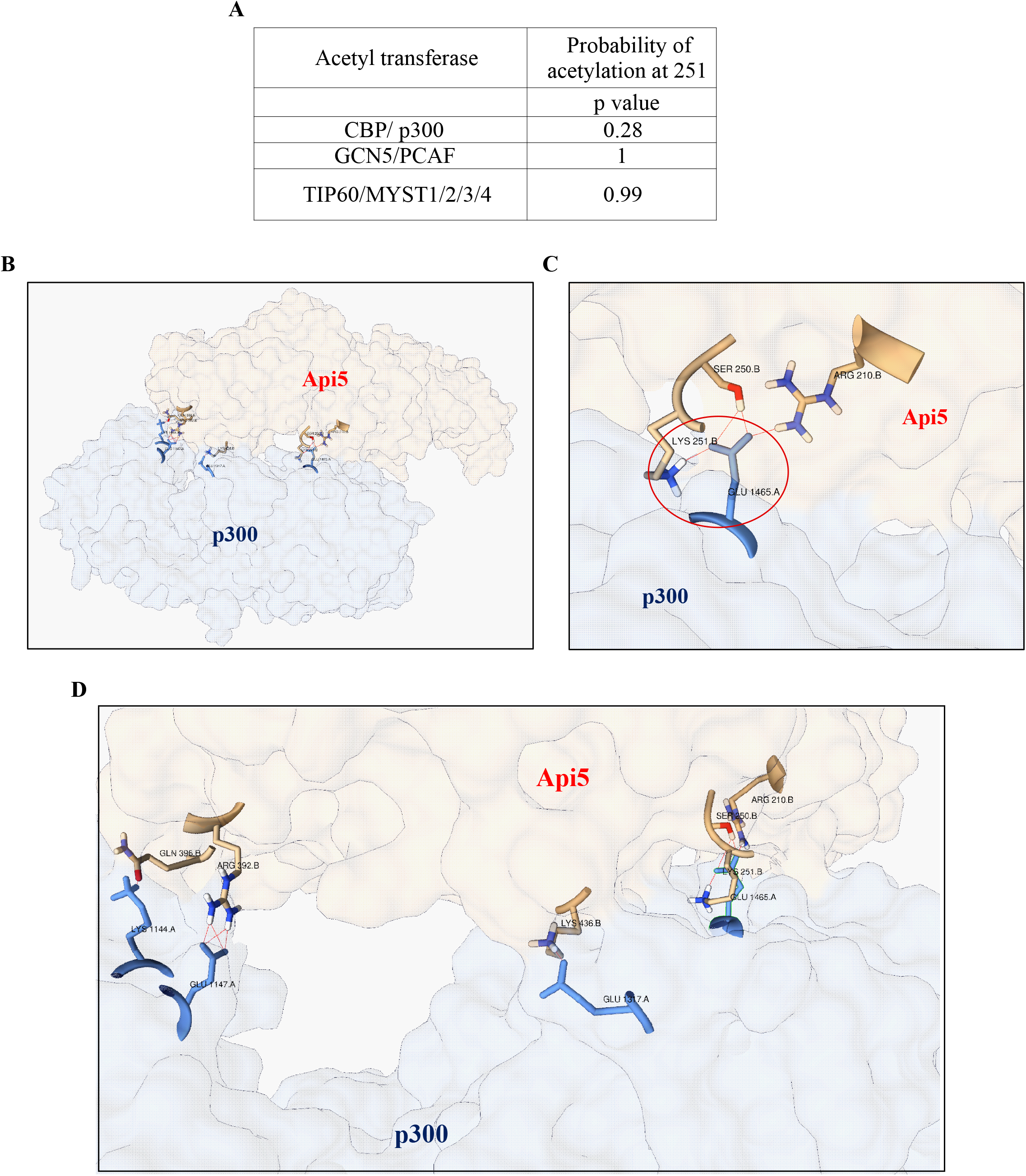
Api5 interacts with p300 *in silico*. A) Various acetyl transferases and their probability to acetylate lysine 251 of Api5 represented as p-value using ASEB webserver, where lower p-value corresponds to a higher probability. B) The lowest-energy docking position of Api5 and p300 analyzed using HADDOCK webserver and output analysed using PIZSA represented as a cartoon diagram. C) Interaction between Lysine 251 of Api5 with p300. D) Various amino acid interactions between Api5 and p300.

### p300 interacts with and regulates Api5

Acetylation of Api5 at K251 is responsible for maintaining its stability [2]. To prove whether p300 histone acetyltransferase can regulate Api5 acetylation and thus stability, cells were treated with different concentrations of a small molecule chemical inhibitor of p300 (p300i) and levels of Api5 were analyzed. Api5 protein expression was reduced upon p300 inhibition (Figure 2A-B) without affecting the transcript expression of Api5 (Figure 2C-D), thus demonstrating that p300 might be regulating Api5 at the post-translational level. In order to acetylate Api5, the interaction between p300 and Api5 is essential. Thus, to find out whether p300 and Api5 physically interact or not, stable cells expressing mCherry-tagged Api5 were immunoprecipitated using GFP- specific antibody (Figure 2E). The presence of p300 histone acetyltransferase in the GFP pull down lysate confirmed the interaction between Api5 with p300. This interaction was further corroborated by reverse immunoprecipitation using p300 antibody and the presence of Api5 was confirmed using GFP specific antibody (Figure 2F). These results conclusively demonstrate p300 to interact with Api5 and thus regulate Api5 protein expression without affecting the transcript levels.

**Figure 2.**
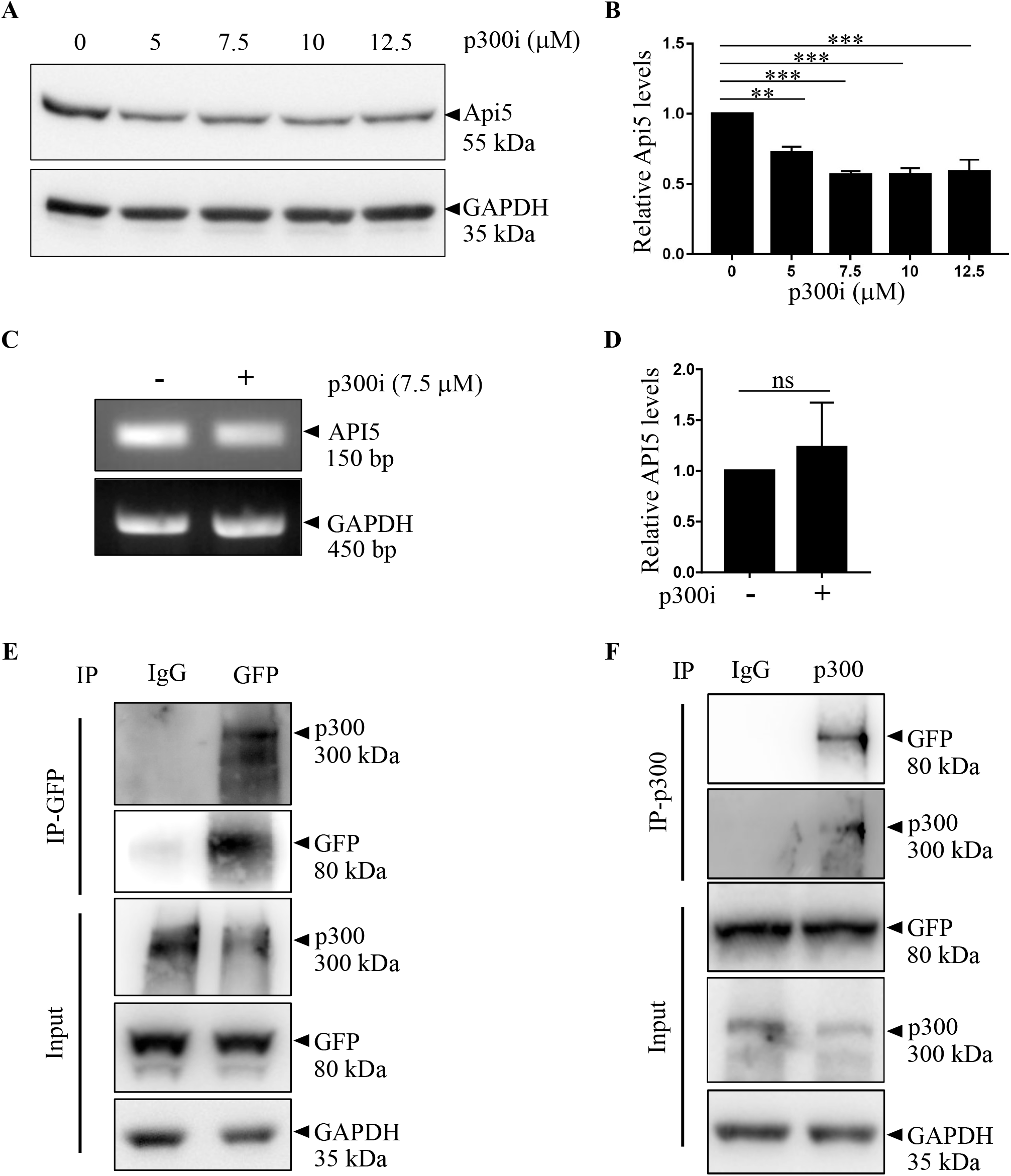
p300 interacts with and regulates Api5. A) MCF-7 cells treated with varying concentrations of p300 inhibitor for 4 hrs were lyzed and Api5 levels were analyzed using western blotting. B) Quantification showing the fold change in Api5 protein levels after normalization with GAPDH. C) API5 transcript levels upon p300 inhibition was analyzed using semi-quantitative PCR. D) quantification of transcript levels after normalizing to GAPDH. MCF-7 cells stably expressing mCherry-Api5 were lyzed and immunoprecipitations were performed using E) GFP and F) p300 specfic antibodies.

### HDAC1 interacts with and regulates levels of Api5

From the previous results, we discovered p300 to be the histone acetyltransferase to regulate Api5 protein levels. Next, we were interested in identifying the de-acetylase enzyme which might be involved in regulating the stability of Api5.

HDAC1 is one of the member of the large family of histone de-acetylases. De-acetylation activity of HDAC1, a class 1 HDAC protein, is coupled with the acetylation activity of p300. Interestingly, HDAC1 has also been reported to be a part of the Api5 interactome [58]. To investigate whether HDAC1 regulates Api5, cells were treated with different concentrations of romidepsin, a HDAC class 1 inhibitor (HDACi), and Api5 protein levels were analyzed. Api5 protein levels increased upon treatment with HDACi suggesting that HDAC1 may be involved in regulating the stability of Api5 (Figure 3A-B). The accumulation of Api5 upon inhibition of HDAC1 could also be the result of increased transcription of Api5. To rule out this possibility, Api5 transcript levels were analyzed upon HDAC1 inhibition. Api5 transcript levels remain unaltered upon romidepsin treatment as shown in Figure 3C-D. These results suggest HDAC1 to play a role in regulating the stability of Api5 at the post translational level. In order to decipher whether HDAC1 interacted with Api5 to bring about the regulation, immunoprecipitation studies were conducted. To demonstrate the interaction of Api5 with HDAC1, whole cell lysates of mCherry-Api5 overexpressing stable cells were immune-precipitated using GFP-specific antibody (Figure 3E). Presence of HDAC1 in the GFP pull down lysates confirmed the interaction between HDAC1 and Api5. This interaction was further corroborated by reverse immuno-precipitation and pull down was performed using HDAC1 specific antibody. Api5 was found to be present in the HDAC1 pull down lysates (Figure 3F). These studies thus validate that HDAC1 interacts with and regulate Api5 at the protein level.

**Figure 3:**
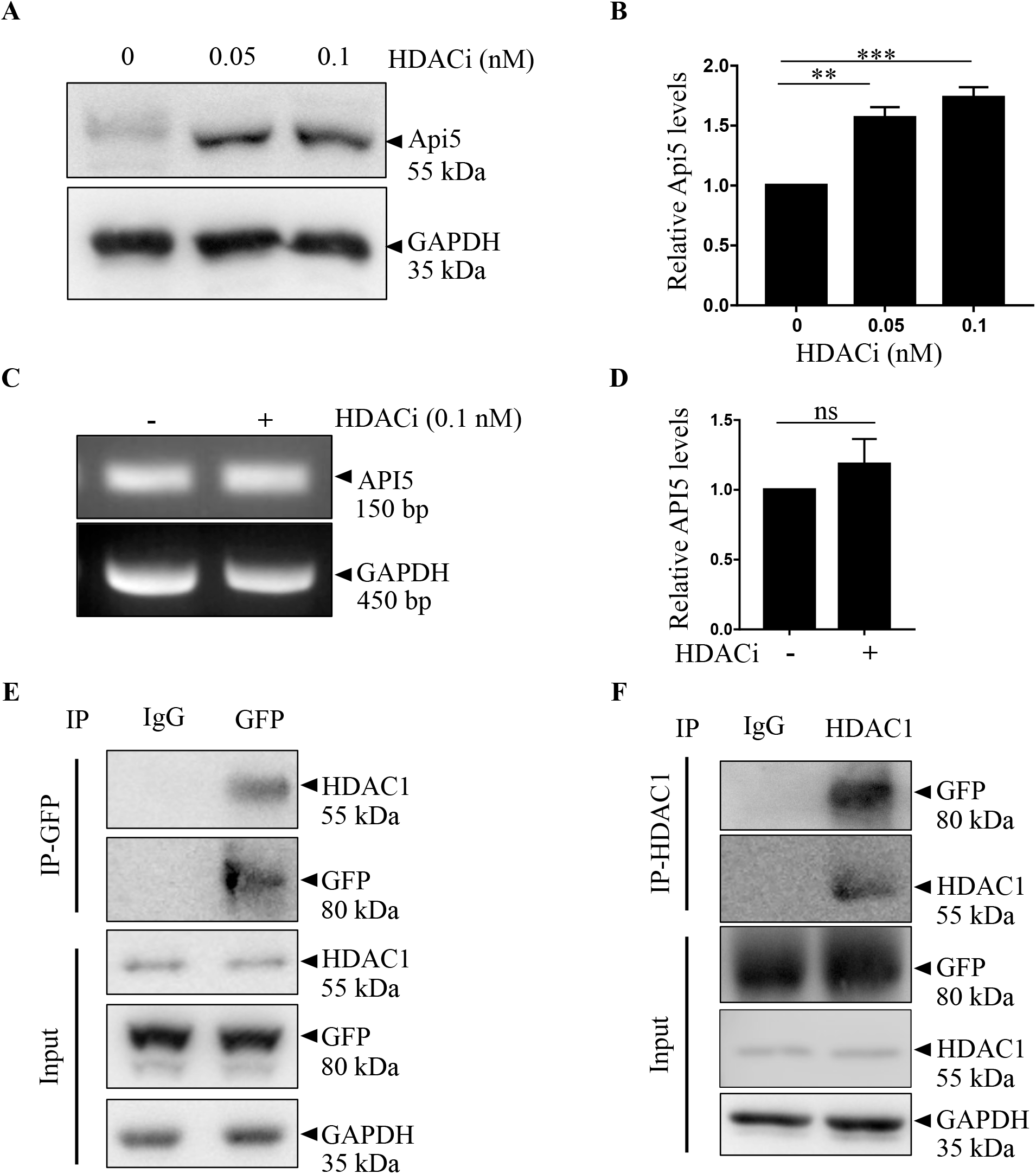
HDAC1 interacts with and regulates Api5. A) MCF-7 cells treated with varying concentrations of the HDAC1 inhibitor, romidepsin for 16 hrs were lyzed and Api5 levels were analyzed using western blotting. B) Quantification showing the fold change in Api5 protein levels after normalization with GAPDH. C) API5 transcript levels upon romidepsin treatment was analyzed using semi-quantitative PCR and D) quantified after normalizing with GAPDH. MCF7 cells stably expressing mCherry-Api5 were lyzed and immunoprecipitation was carried out using E) GFP and F) HDAC1 specfic antibodies.

### Api5 is acetylated by p300 and de-acetylated by HDAC1 at lysine 251 residue

Previous experiments suggested p300 histone acetyltransferase and HDAC1 as the acetylating and de-acetylating enzymes respectively that regulate the levels of Api5. It was hypothesized that these two enzymes might also regulate the dynamics of acetylation and de-acetylation of Api5. Therefore, the acetylation status of Api5 was investigated upon inhibition of both p300 and HDAC1.

To investigate whether Api5 is acetylated by p300, mCherry-Api5 overexpressing stable cells were treated with and without p300i and immunoprecipitation studies were performed. Acetylated Api5 were detected using L-Acetyl lysine specific antibody. Api5 acetylation was observed to be reduced upon inhibition of p300 when compared to the untreated control as shown in Figure 4A. To check for the effect of HDAC1 inhibition using romidepsin on the acetylation status of Api5 similar immunoprecipitation experiments were performed with and without romidepsin. Acetylated levels of Api5 increased upon inhibition of HDAC1 (Figure 4B). This confirms that Api5 undergoes acetylation and de-acetylation by p300 and HDAC1 respectively. Earlier studies by Han and group have shown that Api5 is acetylated at lysine 251 [2]. To identify whether Api5 acetylation and deacetylation by p300 and HDAC1 respectively occurs at the lysine 251 residue, K251 mutants of Api5 were generated using site-directed mutagenesis. mVenus-tagged wild type, K251A (acetylation-deficient mutant), K251R (charge-mimic mutant) and K251Q (constitutive acetylation mimic) mutants of Api5 were ectopically expressed in cells. The expression levels of mVenus, which is a proxy for ectopic expression of Api5 K251 mutants were analyzed in the presence and absence of p300i and romidepsin using GFP-specific antibody. It was observed that levels of wild-type Api5 were reduced while the levels of K251A, K251Q and K251R mutants of Api5 remained unaffected upon p300 inhibition (Figure 4C and D). However, the levels of wild-type Api5 increased upon inhibition of HDAC1 by romidepsin whereas that of the other mutants remained unchanged (Figure 4E and F). Therefore, it was established that p300 acetylates while HDAC1 de-acetylates Api5 at the lysine 251 residue.

**Figure 4:**
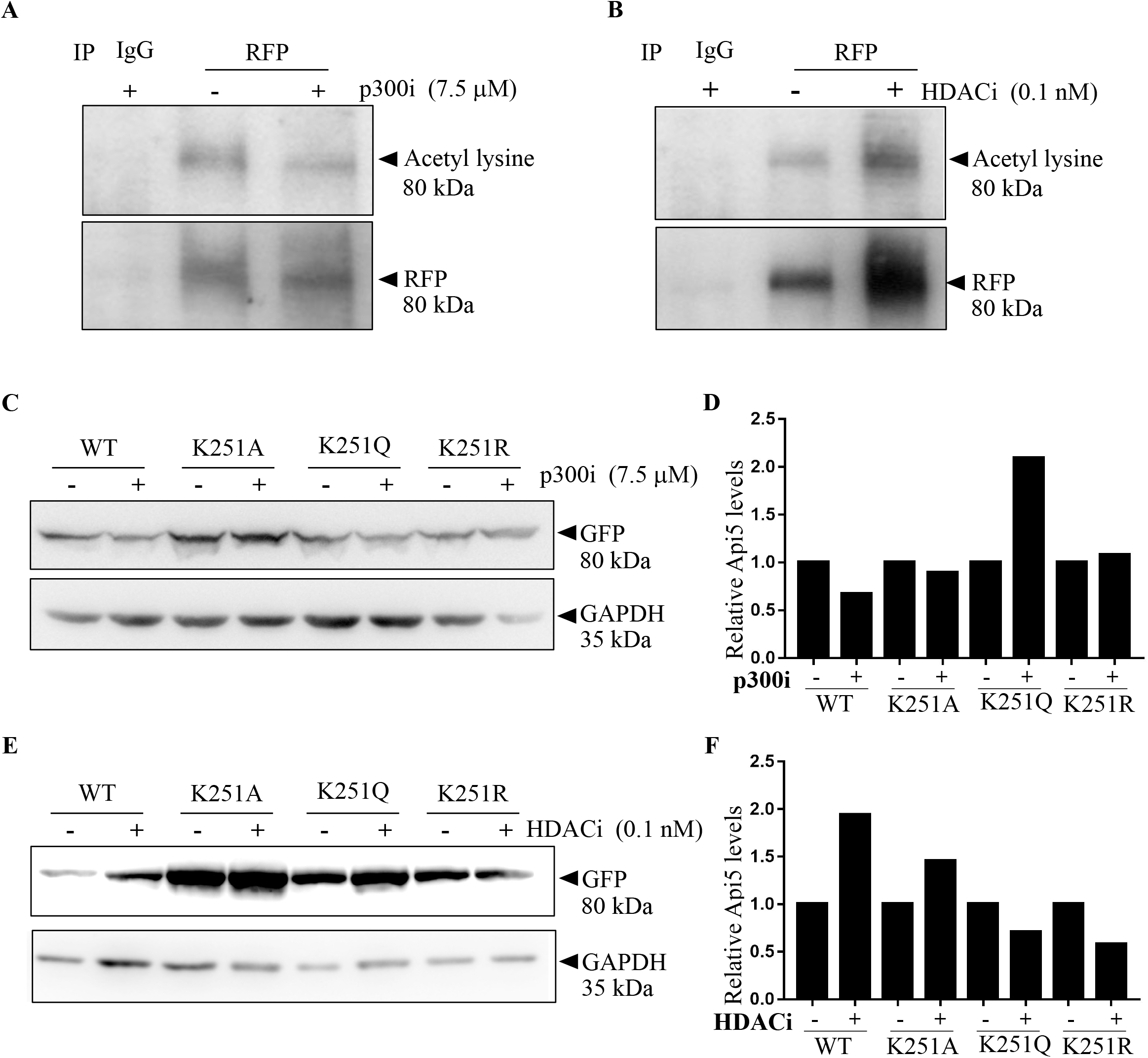
p300 and HDAC1 acetylate and deacetylate Api5 at K251 respectively. mCherry-tagged Api5 over-expressing MCF-7 cells treated with A) 7.5 μM of p300i for 4 hrs and B) 0.1 nM of romidepsin for 16 hrs were lyzed and immunoprecipitation was performed using RFP-specific antibody. Western blotting analysis was performed to ascertain the acetylation status of Api5 using acetyl lysine specific antibody. MCF-7 cells were transfected with K251 mutant constructs and treated with C) 7.5 μM of p300i and E) 0.1 nM of romidepsin post 24 hrs of transfection for 4 and 16 hrs respectively. Western blotting was performed to check for the levels of mutant Api5 using GFP specific antibody. D) and F) Quantification showing the fold change in Api5 protein levels after GAPDH normalization.

### Acetylation of Api5 by p300 is the signal for its nuclear localization while de-acetylation leads to the translocation of Api5 to the cytoplasm

From the previous results, it can be inferred that p300 acetylates Api5 at lysine 251, thereby maintaining its stability. It has earlier been reported that Api5 is a nuclear protein. To investigate the role of acetylation on Api5 function, cells were treated with the acetylation and deacetylation inhibitors and immunofluorescence was performed. In control cells, Api5 was observed to be present in the nucleus while upon inhibition of p300, the percentage of cells showing Api5 to be present in the nucleus was negligible. Moreover, most cells showed nuclear and cytoplasmic localization of Api5 (Figure 5A-B). Interestingly, 24% of cells showed only cytoplasmic presence of Api5 upon p300 inhibition which was not observed in the control cells (Figure 5B). Api5 localization was also investigated upon inhibition of HDAC1. We found that, upon HDAC1 inhibition, majority of the cells (67%) showed both nuclear and cytoplasmic localization of Api5, whereas 29% of cells showed only nuclear localization of Api5 (Figure 5C-D). Only 4% of cells showed cytoplasmic localization of Api5 which is negligible. To follow the compartmentalization switch of Api5, both p300 and HDAC1 were inhibited. The cytoplasmic localization of Api5 that was observed upon p300 inhibition as well as the nuclear localization observed upon HDAC1 inhibition was reduced upon inhibition of both p300 and HDAC1. 92% of cells showed both nuclear and cytoplasmic localization of Api5 (Figure 5E-F). Interestingly it was observed that upon inhibition of HDAC1, 73% of cells showed Api5 nuclear foci while the foci were reduced to 46% when both p300 and HDAC1 inhibitors were added in concert (Figure 5G). Therefore, it was concluded that Api5 acetylation at lysine 251 by p300 stabilizes Api5 inside the nucleus while de-acetylation by HDAC1 at the same lysine residue leads to Api5 to locate to the cytoplasm.

**Figure 5:**
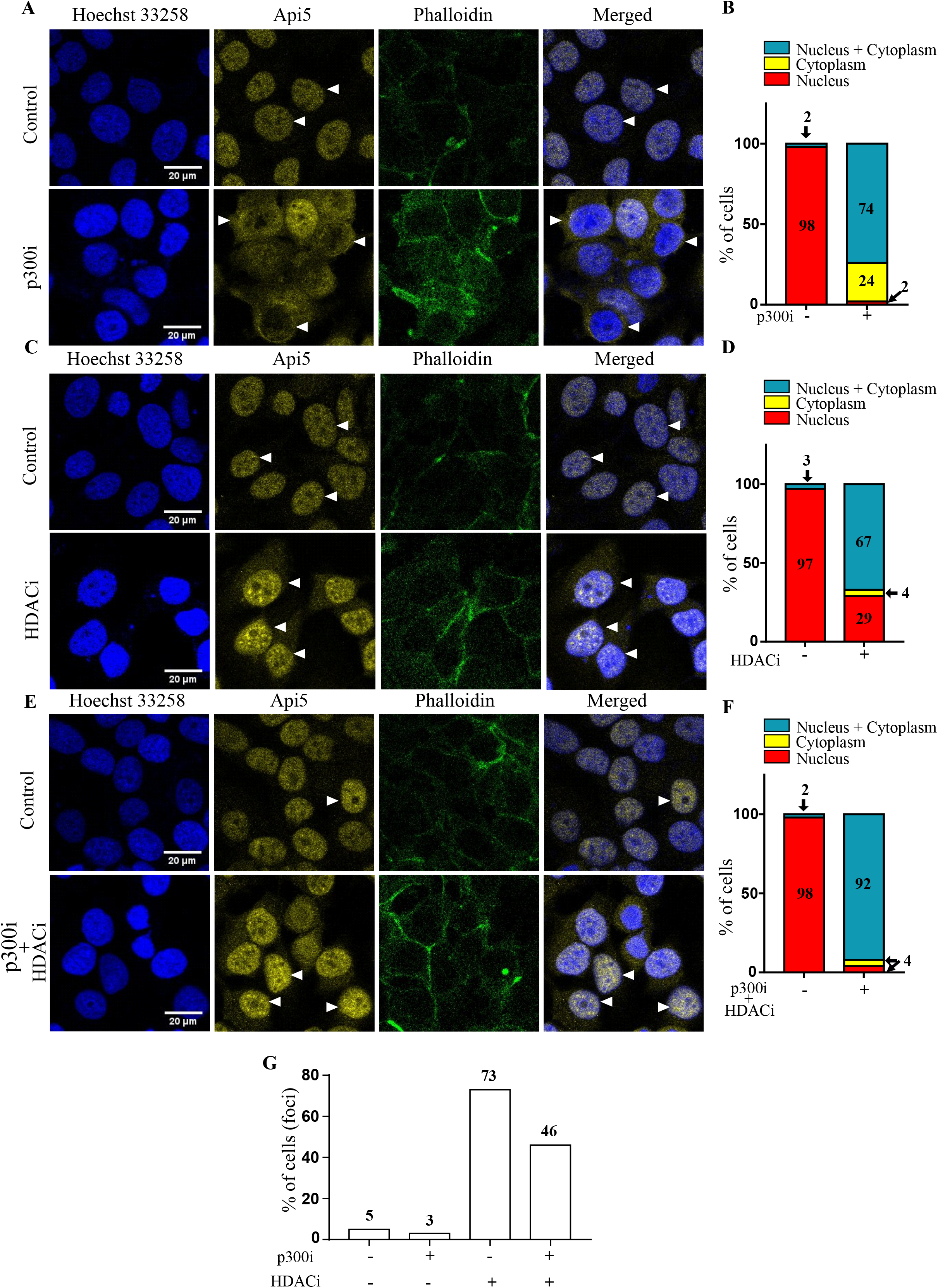
p300 and HDAC1 regulate Api5 localization. MCF-7 cells were treated with A-B) 7.5 μM p300i, C-D) 0.1 nM HDACi and E-F) 7.5 μM p300 and 0.1 nM HDACi for 4 hrs and 16 hrs respectively. Immunofluorescence assay was performed using Api5 specific antibody and cells with different Api5 staining pattern were manually counted and analyzed. G) Percentage of cells showing Api5 foci formation following the different inhibitor treatments.

### Acetylation of Api5 is critical for normal cell cycle progression

Lacazette and group have reported Api5 protein levels to peak during the G1 and S phase of cell cycle while the transcript levels remained unaltered [9]. Our data as well as data from Han’s group [2] have shown acetylation to be responsible for maintaining the stability of Api5. It may be hypothesized that the stability of Api5 during the G1 and S phases of the cell cycle may be due to acetylation of Api5. To demonstrate whether Api5 acetylation affected cell cycle profile, cells were transfected with the K251 mutants of Api5 in an Api5 knock down background and then analyzed. Interestingly, cells transfected with the K251R mutant of Api5 were observed to be arrested at the S phase of the cell cycle (Figure 6A). PCNA, a S-phase cell cycle marker was also found to be upregulated in the Api5-K251R transfected cells (Figure 6B-D) while the other cell cycle markers remained unaffected (Figure 6B, E and F). This result was further corroborated with the observation that the Api5-K251R mutant expressing cells showed higher proliferation when compared to the other Api5 acetylation mutants (Figure 6G). This increase in proliferation may possibly be due to an alteration in the cell cycle functioning resulting from the change in acetylation status of Api5. Therefore it was confirmed that the acetylation status of Api5 also regulated proliferation and cell cycle progression.

**Figure 6:**
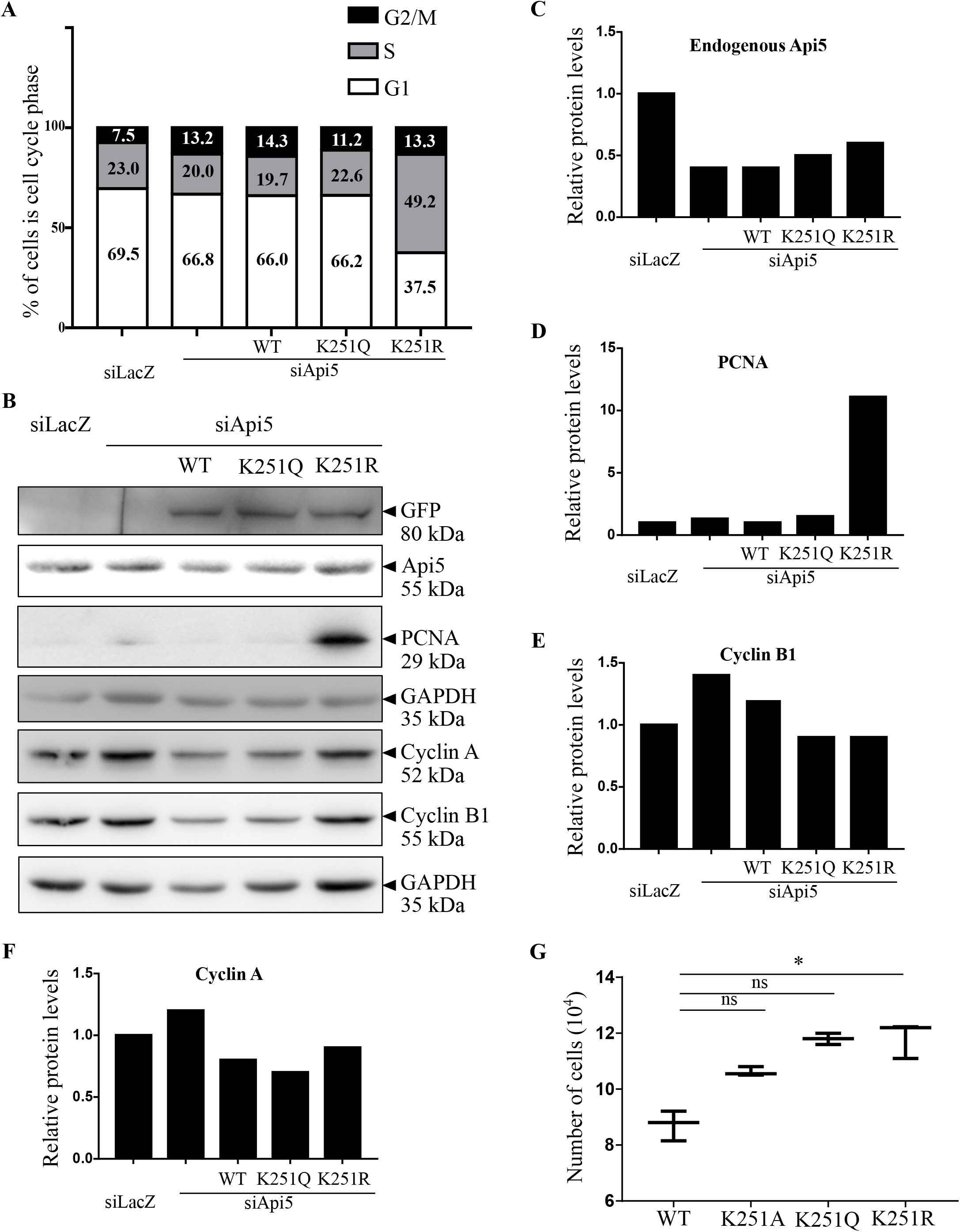
Acetylation status of Api5 affects cell cycle. MCF-7 cells transfected with siRNA-resistant Api5-K251 mutants 24 hrs post Api5 knock down. Cells were fixed in ethanol and stained with propidium iodide and cell cycle profile was monitored. A) Percentage of cells in the different cell cycle phases were analyzed in a flow cytometer and plotted as box plots. B) Western blotting was performed to check the levels of B) Api5, PCNA, Cyclin A and Cyclin B1 and C-F) quantified after normalizing to the loading control GAPDH. G) MCF7 cells transfected with mVenus-tagged Api5 WT and K251 mutants were counted using Sceptar™ to check for cell proliferation.

### AMPK and Akt regulate the p300 mediated stability of Api5

It has already been reported that AMPK can either directly or indirectly regulate G1/S cell cycle transition regulators such as p53 and p21 [47]. AMPK requires the activity of AKT to regulate other cell cycle regulators [59]. It has been previously reported that activated Akt can activate p300 by phosphorylating it at serine 1834 residue [60]. Therefore to discern whether this signaling cascade led to the stability of Api5, both AMPK and Akt were inhibited using small molecule inhibitors. Api5 levels were observed to be reduced upon inhibition of both AMPK (Figure 7A-B) and Akt (Figure 7C-D) independently. It was also observed that upon inhibition of AMPK, Akt phosphorylation at serine 473 decreased which is known to be required to activate p300 (Figure 7E-F). Our experiments suggested that inhibition of AMPK inhibits the activation of Akt. Akt is the kinase that has been reported to phosphorylate and activate p300 histone acetyl transferase. E2F1, a transcription factor is known to be acetylated by p300. This p300-mediated acetylation of E2F1 has been shown to regulate stability of the protein. Reduced levels of E2F1 upon inhibition of AMPK and Akt separately confirms that both AMPK and Akt are involved in regulating the histone acetyl transferase activity of p300 (Figure 7E-I). Inhibition of activation of Akt and reduction in Api5 protein levels upon inhibition of AMPK confirms that p300 mediated stability of Api5 is achieved through the AMPK-Akt pathway as illustrated in Figure 8A.

**Figure 7:**
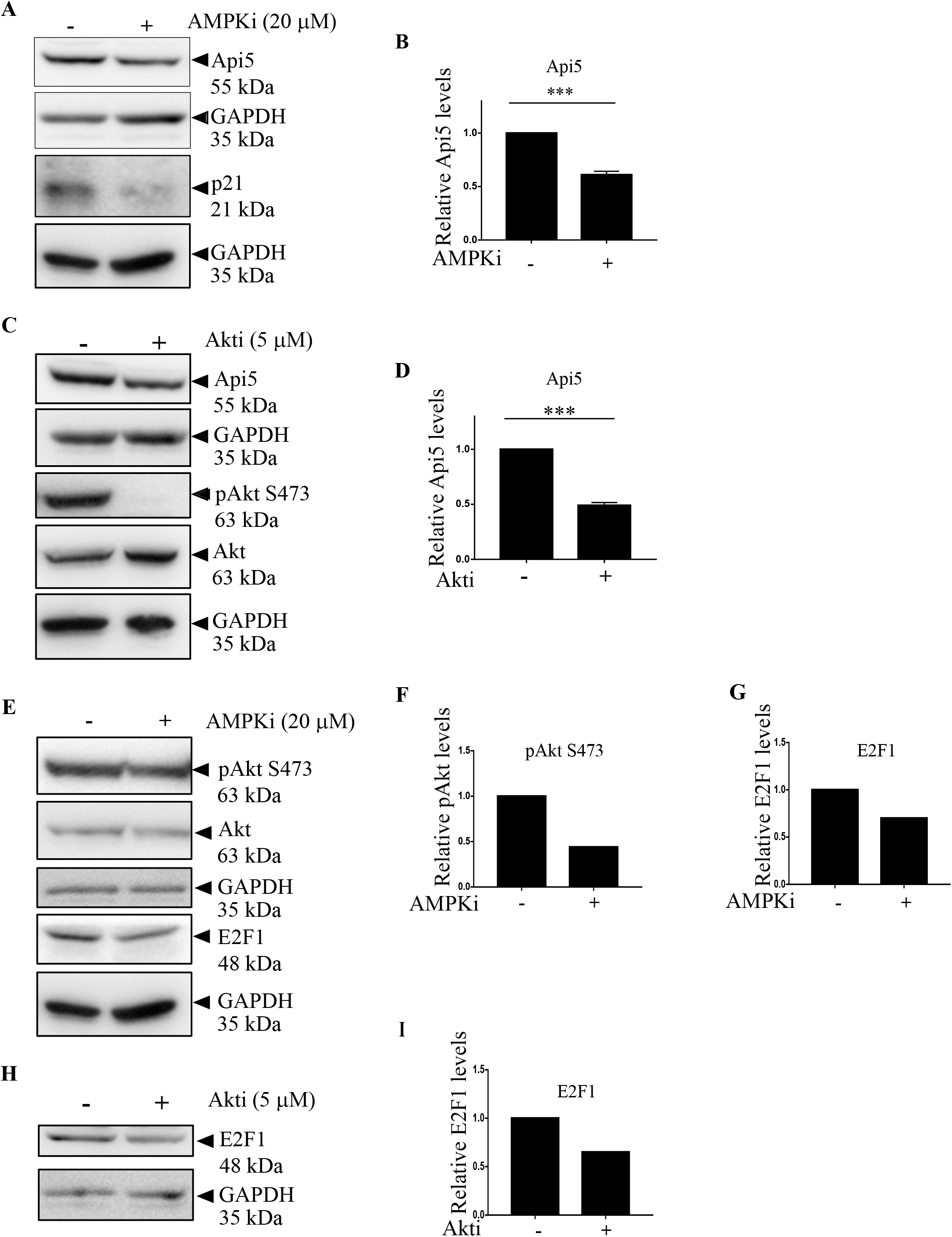
AMPK and AKT regulate Api5 stability. MCF-7 cells were treated with A) 20 μM AMPKi, and C) 5 μM AKTi for 4 hrs and 24 hrs respectively. Immunoblotting analysis was performed using Api5 specific antibody. GAPDH was used as loading control. B and D) Quantification showing the fold change in Api5 levels after normalization to GAPDH. A) p21 and C) pAkt S373 were used as positive control to confirm AMPK and Akt inhibition respectively. E and F) pAkt S473 and E and G) E2F1 levels upon AMPK inhibition. H and I) E2F1 protein levels were analyzed following Akt inhibition using immunoblotting.

**Figure 8:**
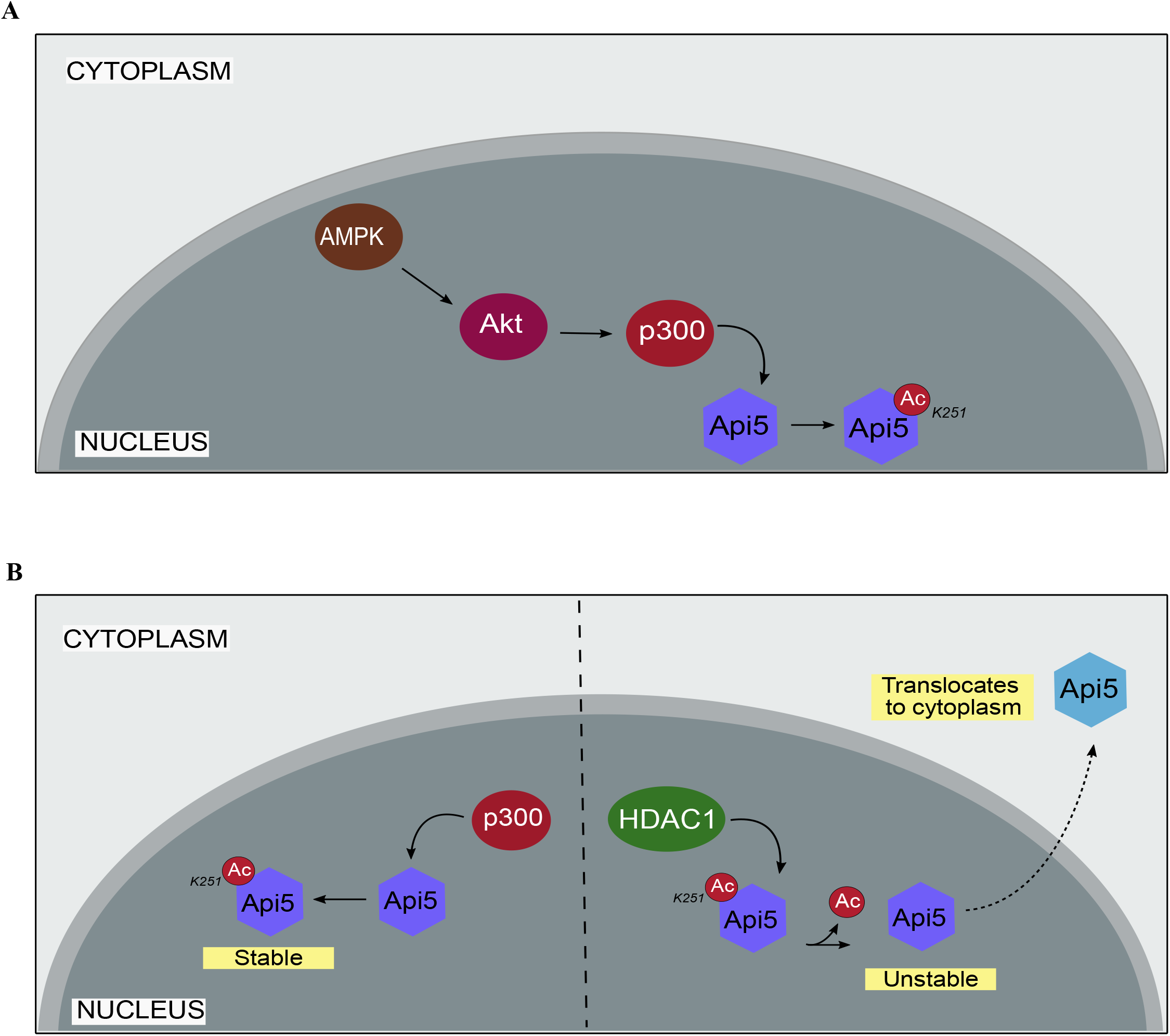
Schematic representation depicting the regulation of Api5 following acetylation and deacetylation. A) Model predicting the regulation of Api5 via the AMPK, Akt and p300 signaling cascade. B) p300 acetylation of Api5 at lysine residue 251 stabilizes the protein in the nucleus; whereas de-acetylation of Api5 at K251 by HDAC1 de-stabilizes the protein which enables it to get translocated to the cytoplasm.

## Discussion

In this study we have shown that p300, a histone acetyltransferase to be responsible for acetylating and maintaining the stability of Api5. p300 is one of the most studied acetyltransferases which not only acetylates histone proteins but also other non-histone proteins too [23]. E2F1 is another known protein that is stabilized upon acetylation by p300 [28]. E2F1 is a transcription factor that plays a critical role during the G1/S transition of the cell cycle [29]. E2F1 levels vary during the different cell cycle phases, similar to that of Api5 [9]. E2F1 protein levels peak during the G1 phase of cell cycle and starts to degrade as the cell crosses the S phase. p53, the master regulator of the cell is also acetylated by the p300 [26]. There are a number of studies that show that upon acetylation, p53 stabilizes and thus, is able to perform its different functions like activation of DNA damage response and repair pathway, cell cycle arrest and apoptosis [42].

Histone deacetylases are one of the members of the deacetylase family of enzymes that remove the acetylation from targeted proteins. Initially HDACs were identified and reported to de-acetylate histone proteins [40]. But extensive studies have shown that HDACs can also de-acetylate non-histone proteins. Both E2F1 and p53, which are acetylated by p300, are also de-acetylated by Class 1 HDACs [29; 61]. E2F1 and p53 upon deacetylation become unstable and lose their activity. HDAC1 mediated de-acetylation of E2F1 also aids in its localization shift from nucleus to cytoplasm.

Earlier studies have shown Api5 acetylation at K251 residue to stabilize Api5 inside the nucleus [2]. However, the enzymes involved in the acetylation process were unknown. Our study, is the first to identify p300 and HDAC1 as the acetylating and de-acetylating enzymes of Api5 respectively, that work in concert, thus leading to the stability or instability of the protein during normal cell functioning. In this study, we report that Api5, a nuclear protein under normal physiological conditions is acetylated by histone acetyltransferase p300. p300 is an essential regulator of Api5 as the acetylated protein is localized in the nucleus while the de-acetylated protein switches its localization to the cytoplasm. As Api5 levels reduce upon inhibition of p300, it suggests that Api5 might be undergoing post-translational degradation in the cytoplasm. As the acetylated Api5 is stable in the nucleus and is not transported to the cytoplasm, it may also be inferred that acetylation prevents Api5 from binding to translocator/transporter proteins that are responsible for its translocation/ transportation from the nuclear compartment to the cytoplasm. However, de-acetylation might be exposing Api5 binding site to transporter protein(s) that enables the protein to translocate to the cytoplasm.

As mentioned earlier, Api5 levels peak during G1 and S phase of cell cycle and is thereby able to play a role in the regulation of cell cycle progression as well as proliferation of cells by possibly functioning as a transcription factor. During G1 and S phases of the cell cycle, Api5 may possibly be interacting with chromatin to transcribe those genes that regulate progression of the cell cycle. Transcription of those genes may also regulate the proliferation of cells. Thus it may be inferred that acetylation of Api5 by p300 possibly provides additional functionality to Api5 that includes transcription of cell cycle regulators.

It has also been reported that Api5 is overexpressed in many cancers. This overexpression of Api5 promotes angiogenesis and metastasis in cancerous cells. Oncogenic properties of tumor promoting proteins can also be considered as an outcome of their stability. In this case, Api5 is being stabilized by the action of p300 histone acetyltransferase. Inhibition of p300 and induction of HDAC1 can reduce the oncogenic potential of Api5. Thus inhibitors of p300 and inducers of HDAC1 have therapeutic implications in the treatment of cancers that have high expression of Api5.

As K251 acetylation of Api5 is the only known post-translational modification which is conserved from protists to mammals [2], suggesting that conservation of this acetylation may play a key role during normal development. AMPK and Akt are important regulators of various cellular processes like cell division, transcription regulation and apoptosis [44; 45; 46] [49; 50]. Reduced levels of Api5 upon inhibition of these key regulators also indicates its role in normal physiological and developmental processes. Although our study suggests that p300 is the acetylating and stabilizing regulator of Api5 while HDAC1 is the de-acetylating and negative regulator of stability, one cannot ignore the possibility of other enzymes too playing a role in the regulation of Api5 stability.

Thus taken together, our data suggests that p300 and HDAC1 are the novel regulators of Api5 which not only interact but also mediate the localization and activity of Api5 by acetylating and de-acetylating at lysine 251 respectively as illustrated in Figure 8B.

## Materials and methods

### Chemicals and antibodies

p300 inhibitor (382113), protease inhibitor cocktail (P2714), dimethyl sulfoxide (DMSO) and propidium iodide (PI) (P4170) were purchased from Sigma-Aldrich and romidepsin from Seleck Biochem. Lipofectamine 2000, Alexa flour 488 goat anti-rabbit and Phalloidin 568 were purchased from Invitrogen. GFP (ab290), RFP (ab62341) and p300 (ab10485) antibodies were obtained from Abcam, HDAC1 (sc-8410) from Santacrutz, Api5 (PAB7951) from Abnova, acetyl lysine (05-515) from Merck Millipore and GAPDH (G9545) from Sigma-Aldrich.

### Plasmids

CSII-EF-MCS plasmid was a gift from Dr. Sourav Banerjee, NBRC, Manesar, India. pCAG-HIVgp and pCMV-VSV-G-RSV-Rev plasmids were purchased from RIKEN BioResource Centre. mVenusC1 was gifted by Jennifer Lippincott-Schwartz, NIH, USA in which Api5 was cloned. The Api5 was made siRNA-resistant and K251 mutants were generated using site directed mutagenesis.

### Cell culture and drug treatments

MCF-7 cells were obtained from the European Collection of Cell Cultures (ECACC). HEK293T cells were a gift from Dr. Jomon Joseph (NCCS, Pune). The cells were maintained in 100 mm dishes (Eppendorf or Corning) and were grown in high glucose Dulbecco’s Modified Eagle Medium (DMEM; Lonza) containing 10% heat inactivated FBS (Invitrogen) and 100 units/mL penicillin-streptomycin (Invitrogen) and incubated at 37°C humidified 5% CO_2_ incubators (Eppendorf or Thermo Scientific).

7 × 10^5^ cell were seeded on 35mm dishes 16 hrs prior to treatment of p300i or HDACi and incubated for 4 hrs and 12 hrs respectively.

MCF-7 cells stably expressing mCherry-tagged Api5 were generated using lentiviral-mediated transduction. Briefly, 7.5 × 10^5^ HEK293T cells were seeded on 35mm dish and transfected with 1 μg of mCherry-CSII-EF-MCS or mCherry-CSII-EF-MCS-Api5 plasmid along with 1 μg of pCAG-HIVgp and 0.5 μg pCMV-VSV-G-RSV-Rev packaging plasmids using Lipofectamin 2000. 1 ml DMEM containing 30% FBS was added to the cells 24 hours post transfection. 5 × 10^5^ MCF-7 cells were seeded on a 35 mm dish for transduction. Viral supernatant was collected 48 hours post transfection and filtered through a 0.45 μm filter to get rid of the cell debris. Filtered viral supernatant containing media along with 1 ml fresh media was added to the MCF-7 cells. 4 μg polybrene was added to the cells to increase the transduction efficiency. Cells were replenished with fresh medium 48 hours post transduction. Transduced MCF-7 cells were passaged and used for further experiments.

For transfections, 6 × 10^5^ cells were seeded in 35mm dishes and incubated at 37°C overnight. siRNA duplexes targeting Api5 and LacZ were purchased from Dharmacon (Thermo Scientific). Sense sequences of the siRNA are: Api5, 5′-GACCUAGAACAGACCUUCAUU-3′, LacZ, 5′-CGUACGCGGAAUA CUUCGAdTdT-3′. siRNA against LacZ and Api5 were transfected as described earlier [62]. Api5 K251 mutants which were generated by site directed mutagenesis were transfected 24 hrs post knockdown of Api5.

### Immunoblotting and Immunoprecipitation

mCherry-tagged Api5 over-expressing stable cells with or without treatments were lyzed in cell lysis buffer containing 50 mM Tris-HCl, pH 7.4, 0.1% Triton X-100, 5 mM EDTA, 250 mM NaCl, 50 mM NaF, 0.1 mM Na_3_VO_4_ and protease inhibitors. Immunoprecipitations were performed as described [63] using GFP, RFP, p300, or HDAC1 specific antibodies. Western blotting was performed with Api5, GFP, acetyl lysine and GAPDH antibodies as previously mentioned [62].

### Semi-quantitative PCR

RNA extraction and cDNA synthesis of p300i and HDACi treated cells were performed as described previously [64]. Semi-quantitative PCR was performed using API5 specific forward (5’-CGAGTGGCAGATATACTAACGC-3’) and reverse (5’-TCCTCTCCTTGAAGTATTTGGC-3’) primers. GAPDH was used as endogenous control and was amplified using forward (5’-ACCACAGTCCATGCCATCAC-3’) and reverse (5’-TCCACACCCTGTTGCTGTA-3’) primers. The following PCR cycle was used for the amplification: 95°C for 60 sec, 58°C for 45 sec, 72°C for 60 sec and final extension for 3 min. The experiments were repeated three times to confirm API5 levels

### Immunofluorescence

6 × 10^5^ cells were seeded on coverslips placed in 35mm dishes. After treatment of p300i or HDACi, cells were fixed in 4% paraformaldehyde and permeabilized using 0.5% TritonX after PBS washes. After PBS-Glycine wash, cells were blocked using 10% FBS for 1 hr and incubated overnight with Api5 antibody at 4°C. Cells were incubated with secondary antibodies conjugated with Alexa Flour for 1 hr at room temperature along with phalloidin. Nuclei was stained using Hoechst 33258 for 5 mins and washed twice using PBS. The cells were mounted using mounting media and imaged under 63X oil immersion objective of the SP8 confocal microscope (Leica, Germany).

### Cell proliferation assay

6 × 10^5^ MCF-7 cells were seeded in 35mm dishes and incubated at 37°C overnight. mVenus-tagged WT and K251 mutants of Api5 plasmids were transfected as mentioned before. 48 hrs post transfection, cells were trypsinized and pelleted down by centrifugation at 1000 rpm for 5 mins. Cell pellet was suspended in 1X PBS. 1:10 dilution of the suspended cell pellet was prepared in 1X PBS and the cell count was measured using 60 μm sensor connected to Sceptar™ hand held automated cell counter (Millipore, PHCC00000).

### Flow cytometry

MCF-7 cells after transfection were trypsinized using 1X trypsin and collected in media. After pelleting, the cells were washed with cold PBS twice to remove excess media. The cells were then fixed in 70% ethanol overnight at 4°C and stained with 1 mg/ml Propidium Iodide in the presence of RNase A (20 mg/ml) at 37°C for 1 hr. Cell cycle profile was analyzed on BD Accuri™ C6 flow cytometer (BD Biosciences).

### Statistical analysis

Densitometry analysis of western blots were performed using ImageJ software. Results were analyzed from a minimumm of three independent experiments and plotted using GraphPad Prism (GraphPad Software, La Jolla, CA, USA). Student’s t-test and Mann Whitney U test was used to test the significance in difference of Api5 levels between two samples and One-way Anova and Dunnett’s multiple comparisons test for more than two samples. p value > 0.05 was considered non-significant. *, ** and *** correspond to p<0.05, p<0.001, and p<0.001 respectively.

## Conflict of Interest

The authors declare that they have no competing interests.

## Author Contributions

V.K.S and M.L conceived and conceptualized the project. V.K.S designed, performed and analyzed the experiments. V.K.S and M.L wrote the paper.

## Funding

This study is supported by a grant from Science and Engineering Research Board (SERB), Govt. of India (EMR/2016/001974) and partly by IISER, Pune Core funding. V.K.S was funded by IISER Pune Core funding.

## Acknowledgments

We thank Drs Richa Rikhy, Nagaraj Balasubramanium (IISER Pune, India) and Manas Kumar Santra (NCCS, Pune, India) for their useful suggestions. We also thank Dr. Krishanpal Karmodiya for the HDAC inhibitor and Dr. Gayathri Panangat as well as Sanjana Nair (IISER Pune, India) for help with the *in-silico* analysis. The authors acknowledge the IISER Pune Microscopy Facility for access to equipment and infrastructure. We thank Lahiri Lab members Rintu Umesh for help with stable cell line preparation and Aishwarya Venkataravi for designing the schematics in Figure 8. We also thank Lahiri lab members for helpful comments and discussions

## Notes

### Competing Interest Statement

The authors have declared no competing interest.

## References

[1] M. Tewari, M. Yu, B. Ross, C. Dean, A. Giordano, and R. Rubin, AAC-11, a novel cDNA that inhibits apoptosis after growth factor withdrawal. Cancer research 57 (1997) 4063–9.

[2] B.G. Han, K.H. Kim, S.J. Lee, K.C. Jeong, J.W. Cho, K.H. Noh, T.W. Kim, S.J. Kim, H.J. Yoon, S.W. Suh, S. Lee, and B.I. Lee, Helical repeat structure of apoptosis inhibitor 5 reveals protein-protein interaction modules. The Journal of biological chemistry 287 (2012) 10727–37.

[3] L. Van den Berghe, H. Laurell, I. Huez, C. Zanibellato, H. Prats, and B. Bugler, FIF [fibroblast growth factor-2 (FGF-2)-interacting-factor], a nuclear putatively antiapoptotic factor, interacts specifically with FGF-2. Molecular endocrinology 14 (2000) 1709–24.

[4] P. Rigou, V. Piddubnyak, A. Faye, J.C. Rain, L. Michel, F. Calvo, and J.L. Poyet, The antiapoptotic protein AAC-11 interacts with and regulates Acinus-mediated DNA fragmentation. The EMBO journal 28 (2009) 1576–88.

[5] E.J. Morris, W.A. Michaud, J.Y. Ji, N.S. Moon, J.W. Rocco, and N.J. Dyson, Functional identification of Api5 as a suppressor of E2F-dependent apoptosis in vivo. PLoS genetics 2 (2006) e196.

[6] K.H. Noh, S.H. Kim, J.H. Kim, K.H. Song, Y.H. Lee, T.H. Kang, H.D. Han, A.K. Sood, J. Ng, K. Kim, C.H. Sonn, V. Kumar, C. Yee, K.M. Lee, and T.W. Kim, API5 confers tumoral immune escape through FGF2-dependent cell survival pathway. Cancer research 74 (2014) 3556–66.

[7] A.K. Mayank, S. Sharma, H. Nailwal, and S.K. Lal, Nucleoprotein of influenza A virus negatively impacts antiapoptotic protein API5 to enhance E2F1-dependent apoptosis and virus replication. Cell death & disease 6 (2015) e2018.

[8] G. Imre, J. Berthelet, J. Heering, S. Kehrloesser, I.M. Melzer, B.I. Lee, B. Thiede, V. Dotsch, and K. Rajalingam, Apoptosis inhibitor 5 is an endogenous inhibitor of caspase-2. EMBO reports 18 (2017) 733–744.

[9] M. Garcia-Jove Navarro, C. Basset, T. Arcondeguy, C. Touriol, G. Perez, H. Prats, and E. Lacazette, Api5 contributes to E2F1 control of the G1/S cell cycle phase transition. PloS one 8 (2013) e71443.

[10] L. Koci, K. Chlebova, M. Hyzdalova, J. Hofmanova, M. Jira, P. Kysela, A. Kozubik, Z. Kala, and P. Krejci, Apoptosis inhibitor 5 (API-5; AAC-11; FIF) is upregulated in human carcinomas in vivo. Oncology letters 3 (2012) 913–916.

[11] H. Cho, J.Y. Chung, K.H. Song, K.H. Noh, B.W. Kim, E.J. Chung, K. Ylaya, J.H. Kim, T.W. Kim, S.M. Hewitt, and J.H. Kim, Apoptosis inhibitor-5 overexpression is associated with tumor progression and poor prognosis in patients with cervical cancer. BMC cancer 14 (2014) 545.

[12] K.H. Song, H. Cho, S. Kim, H.J. Lee, S.J. Oh, S.R. Woo, S.O. Hong, H.S. Jang, K.H. Noh, C.H. Choi, J.Y. Chung, S.M. Hewitt, J.H. Kim, M. Son, S.H. Kim, B.I. Lee, H.C. Park, Y.K. Bae, and T.W. Kim, API5 confers cancer stem cell-like properties through the FGF2-NANOG axis. Oncogenesis 6 (2017) e285.

[13] K. Lawrenson, P. Mhawech-Fauceglia, J. Worthington, T.J. Spindler, D. O’Brien, J.M. Lee, G. Spain, M. Sharifian, G. Wang, K.M. Darcy, T. Pejovic, H. Sowter, J.F. Timms, and S.A. Gayther, Identification of novel candidate biomarkers of epithelial ovarian cancer by profiling the secretomes of three-dimensional genetic models of ovarian carcinogenesis. International journal of cancer 137 (2015) 1806–17.

[14] C. Basset, F. Bonnet-Magnaval, M.G. Navarro, C. Touriol, M. Courtade, H. Prats, B. Garmy-Susini, and E. Lacazette, Api5 a new cofactor of estrogen receptor alpha involved in breast cancer outcome. Oncotarget 8 (2017) 52511–52526.

[15] J. Neuweiler, J. Venturini, and I. Balazs, Properties of a highly polymorphic locus (D2S92) located in the telomeric region of chromosome 2. Nucleic acids research 19 (1991) 6971.

[16] M.P. Jansen, J.A. Foekens, I.L. van Staveren, M.M. Dirkzwager-Kiel, K. Ritstier, M.P. Look, M.E. Meijer-van Gelder, A.M. Sieuwerts, H. Portengen, L.C. Dorssers, J.G. Klijn, and E.M. Berns, Molecular classification of tamoxifen-resistant breast carcinomas by gene expression profiling. Journal of clinical oncology: official journal of the American Society of Clinical Oncology 23 (2005) 732–40.

[17] P. Ramdas, M. Rajihuzzaman, S.D. Veerasenan, K.R. Selvaduray, K. Nesaretnam, and A.K. Radhakrishnan, Tocotrienol-treated MCF-7 human breast cancer cells show down-regulation of API5 and up-regulation of MIG6 genes. Cancer genomics & proteomics 8 (2011) 19–31.

[18] R.H. Giles, D.J. Peters, and M.H. Breuning, Conjunction dysfunction: CBP/p300 in human disease. Trends in genetics: TIG 14 (1998) 178–83.

[19] A. Giordano, and M.L. Avantaggiati, p300 and CBP: partners for life and death. Journal of cellular physiology 181 (1999) 218–30.

[20] R.H. Goodman, and S. Smolik, CBP/p300 in cell growth, transformation, and development. Genes & development 14 (2000) 1553–77.

[21] H.M. Chan, and N.B. La Thangue, p300/CBP proteins: HATs for transcriptional bridges and scaffolds. Journal of cell science 114 (2001) 2363–73.

[22] A.J. Bannister, and T. Kouzarides, The CBP co-activator is a histone acetyltransferase. Nature 384 (1996) 641–3.

[23] V.V. Ogryzko, R.L. Schiltz, V. Russanova, B.H. Howard, and Y. Nakatani, The transcriptional coactivators p300 and CBP are histone acetyltransferases. Cell 87 (1996) 953–9.

[24] W. Gu, and R.G. Roeder, Activation of p53 sequence-specific DNA binding by acetylation of the p53 C-terminal domain. Cell 90 (1997) 595–606.

[25] J. Boyes, P. Byfield, Y. Nakatani, and V. Ogryzko, Regulation of activity of the transcription factor GATA-1 by acetylation. Nature 396 (1998) 594–8.

[26] K. Sakaguchi, J.E. Herrera, S. Saito, T. Miki, M. Bustin, A. Vassilev, C.W. Anderson, and E. Appella, DNA damage activates p53 through a phosphorylation-acetylation cascade. Genes & development 12 (1998) 2831–41.

[27] A.J. Bannister, E.A. Miska, D. Gorlich, and T. Kouzarides, Acetylation of importin-alpha nuclear import factors by CBP/p300. Current biology: CB 10 (2000) 467–70.

[28] M.A. Martinez-Balbas, U.M. Bauer, S.J. Nielsen, A. Brehm, and T. Kouzarides, Regulation of E2F1 activity by acetylation. The EMBO journal 19 (2000) 662–71.

[29] G. Marzio, C. Wagener, M.I. Gutierrez, P. Cartwright, K. Helin, and M. Giacca, E2F family members are differentially regulated by reversible acetylation. The Journal of biological chemistry 275 (2000) 10887–92.

[30] M. Yamamoto, K. Morita, Y. Tomita, K. Tsuji, K. Kawamura, and H. Maeda, Effect of facial affect stimuli on auditory and visual P300 in healthy subjects. The Kurume medical journal 47 (2000) 285–90.

[31] W.Y. Chan, and T.B. Ng, Comparison of the embryotoxic effects of saporin, agrostin (type 1 ribosome-inactivating proteins) and ricin (a type 2 ribosome-inactivating protein). Pharmacology & toxicology 88 (2001) 300–3.

[32] A. Polesskaya, I. Naguibneva, A. Duquet, E. Bengal, P. Robin, and A. Harel-Bellan, Interaction between acetylated MyoD and the bromodomain of CBP and/or p300. Molecular and cellular biology 21 (2001) 5312–20.

[33] A. Costanzo, P. Merlo, N. Pediconi, M. Fulco, V. Sartorelli, P.A. Cole, G. Fontemaggi, M. Fanciulli, L. Schiltz, G. Blandino, C. Balsano, and M. Levrero, DNA damage-dependent acetylation of p73 dictates the selective activation of apoptotic target genes. Molecular cell 9 (2002) 175–86.

[34] S. Ait-Si-Ali, A. Polesskaya, S. Filleur, R. Ferreira, A. Duquet, P. Robin, A. Vervish, D. Trouche, F. Cabon, and A. Harel-Bellan, CBP/p300 histone acetyl-transferase activity is important for the G1/S transition. Oncogene 19 (2000) 2430–7.

[35] J.A. Howe, J.S. Mymryk, C. Egan, P.E. Branton, and S.T. Bayley, Retinoblastoma growth suppressor and a 300-kDa protein appear to regulate cellular DNA synthesis. Proceedings of the National Academy of Sciences of the United States of America 87 (1990) 5883–7.

[36] S.A. Gayther, S.J. Batley, L. Linger, A. Bannister, K. Thorpe, S.F. Chin, Y. Daigo, P. Russell, A. Wilson, H.M. Sowter, J.D. Delhanty, B.A. Ponder, T. Kouzarides, and C. Caldas, Mutations truncating the EP300 acetylase in human cancers. Nature genetics 24 (2000) 300–3.

[37] E.J. Bryan, V.J. Jokubaitis, N.L. Chamberlain, S.W. Baxter, E. Dawson, D.Y. Choong, and I.G. Campbell, Mutation analysis of EP300 in colon, breast and ovarian carcinomas. International journal of cancer 102 (2002) 137–41.

[38] G.W. Tillinghast, J. Partee, P. Albert, J.M. Kelley, K.H. Burtow, and K. Kelly, Analysis of genetic stability at the EP300 and CREBBP loci in a panel of cancer cell lines. Genes, chromosomes & cancer 37 (2003) 121–31.

[39] N. Koshiishi, J.M. Chong, T. Fukasawa, R. Ikeno, Y. Hayashi, N. Funata, H. Nagai, M. Miyaki, Y. Matsumoto, and M. Fukayama, p300 gene alterations in intestinal and diffuse types of gastric carcinoma. Gastric cancer: official journal of the International Gastric Cancer Association and the Japanese Gastric Cancer Association 7 (2004) 85–90.

[40] A. Inoue, and D. Fujimoto, Enzymatic deacetylation of histone. Biochemical and biophysical research communications 36 (1969) 146–50.

[41] R. Marmorstein, and M.M. Zhou, Writers and readers of histone acetylation: structure, mechanism, and inhibition. Cold Spring Harbor perspectives in biology 6 (2014) a018762.

[42] J. Luo, F. Su, D. Chen, A. Shiloh, and W. Gu, Deacetylation of p53 modulates its effect on cell growth and apoptosis. Nature 408 (2000) 377–81.

[43] C. Hubbert, A. Guardiola, R. Shao, Y. Kawaguchi, A. Ito, A. Nixon, M. Yoshida, X.F. Wang, and T.P. Yao, HDAC6 is a microtubule-associated deacetylase. Nature 417 (2002) 455–8.

[44] M.C. Towler, and D.G. Hardie, AMP-activated protein kinase in metabolic control and insulin signaling. Circulation research 100 (2007) 328–41.

[45] S. Fogarty, and D.G. Hardie, Development of protein kinase activators: AMPK as a target in metabolic disorders and cancer. Biochimica et biophysica acta 1804 (2010) 581–91.

[46] N. Musi, and L.J. Goodyear, Targeting the AMP-activated protein kinase for the treatment of type 2 diabetes. Current drug targets. Immune, endocrine and metabolic disorders 2 (2002) 119–27.

[47] D. Kong, Y. Dagon, J.N. Campbell, Y. Guo, Z. Yang, X. Yi, P. Aryal, K. Wellenstein, B.B. Kahn, B.L. Sabatini, and B.B. Lowell, A Postsynaptic AMPK-->p21-Activated Kinase Pathway Drives Fasting-Induced Synaptic Plasticity in AgRP Neurons. Neuron 91 (2016) 25–33.

[48] G. Rehman, A. Shehzad, A.L. Khan, and M. Hamayun, Role of AMP-activated protein kinase in cancer therapy. Archiv der Pharmazie 347 (2014) 457–68.

[49] J. Jin, L. Jin, S.W. Lim, and C.W. Yang, Klotho Deficiency Aggravates Tacrolimus-Induced Renal Injury via the Phosphatidylinositol 3-Kinase-Akt-Forkhead Box Protein O Pathway. American journal of nephrology 43 (2016) 357–65.

[50] Z.H. Ouyang, W.J. Wang, Y.G. Yan, B. Wang, and G.H. Lv, The PI3K/Akt pathway: a critical player in intervertebral disc degeneration. Oncotarget 8 (2017) 57870–57881.

[51] F. Chang, J.T. Lee, P.M. Navolanic, L.S. Steelman, J.G. Shelton, W.L. Blalock, R.A. Franklin, and J.A. McCubrey, Involvement of PI3K/Akt pathway in cell cycle progression, apoptosis, and neoplastic transformation: a target for cancer chemotherapy. Leukemia 17 (2003) 590–603.

[52] P. Liu, H. Cheng, T.M. Roberts, and J.J. Zhao, Targeting the phosphoinositide 3-kinase pathway in cancer. Nature reviews. Drug discovery 8 (2009) 627–44.

[53] L. Wang, Y. Du, M. Lu, and T. Li, ASEB: a web server for KAT-specific acetylation site prediction. Nucleic acids research 40 (2012) W376–9.

[54] G.C.P. van Zundert, J. Rodrigues, M. Trellet, C. Schmitz, P.L. Kastritis, E. Karaca, A.S.J. Melquiond, M. van Dijk, S.J. de Vries, and A. Bonvin, The HADDOCK2.2 Web Server: User-Friendly Integrative Modeling of Biomolecular Complexes. Journal of molecular biology 428 (2016) 720–725.

[55] B.M. Dancy, and P.A. Cole, Protein lysine acetylation by p300/CBP. Chemical reviews 115 (2015) 2419–52.

[56] A.S. Dhawanjewar, A.A. Roy, and M.S. Madhusudhan, A knowledge-based scoring function to assess quaternary associations of proteins. Bioinformatics 36 (2020) 3739–3748.

[57] A.A. Roy, A.S. Dhawanjewar, P. Sharma, G. Singh, and M.S. Madhusudhan, Protein Interaction Z Score Assessment (PIZSA): an empirical scoring scheme for evaluation of protein-protein interactions. Nucleic acids research 47 (2019) W331–W337.

[58] S.M. Bong, S.H. Bae, B. Song, H. Gwak, S.W. Yang, S. Kim, S. Nam, K. Rajalingam, S.J. Oh, T.W. Kim, S. Park, H. Jang, and B.I. Lee, Regulation of mRNA export through API5 and nuclear FGF2 interaction. Nucleic acids research 48 (2020) 6340–6352.

[59] F. Han, C.F. Li, Z. Cai, X. Zhang, G. Jin, W.N. Zhang, C. Xu, C.Y. Wang, J. Morrow, S. Zhang, D. Xu, G. Wang, and H.K. Lin, The critical role of AMPK in driving Akt activation under stress, tumorigenesis and drug resistance. Nature communications 9 (2018) 4728.

[60] W.C. Huang, and C.C. Chen, Akt phosphorylation of p300 at Ser-1834 is essential for its histone acetyltransferase and transcriptional activity. Molecular and cellular biology 25 (2005) 6592–602.

[61] K.L. Harms, and X. Chen, Histone deacetylase 2 modulates p53 transcriptional activities through regulation of p53-DNA binding activity. Cancer research 67 (2007) 3145–52.

[62] S. Bodakuntla, A.V. Libi, S. Sural, P. Trivedi, and M. Lahiri, N-nitroso-N-ethylurea activates DNA damage surveillance pathways and induces transformation in mammalian cells. BMC cancer 14 (2014) 287.

[63] M.K. Santra, N. Wajapeyee, and M.R. Green, F-box protein FBXO31 mediates cyclin D1 degradation to induce G1 arrest after DNA damage. Nature 459 (2009) 722–5.

[64] V.L. Anandi, K.A. Ashiq, K. Nitheesh, and M. Lahiri, Platelet-activating factor promotes motility in breast cancer cells and disrupts non-transformed breast acinar structures. Oncology reports 35 (2016) 179–88.

